# Neighborhood environment, social cohesion, and epigenetic aging

**DOI:** 10.1101/2020.10.19.345777

**Authors:** Chantel L. Martin, Cavin K. Ward-Caviness, Radhika Dhingra, Tarek M. Zikry, Sandro Galea, Derek E. Wildman, Karestan C. Koenen, Monica Uddin, Allison E Aiello

## Abstract

Living in adverse neighborhood environments have been linked to increased risk of aging-related diseases and mortality; however, the biological mechanisms explaining this observation remain poorly understood. DNA methylation (DNAm), a proposed biomarker of biological aging responsive to environmental stressors, offers promising insight into molecular pathways. We examined associations of three measures of neighborhood conditions (poverty, quality, and social cohesion) with three different epigenetic clocks (Horvath, Hannum, and Levine) using data from the Detroit Neighborhood Health Study (n=158). Using linear regression models, we evaluated associations in the total sample and stratified by gender and social cohesion. Differential effects by gender were found between men and women. Neighborhood poverty was associated with PhenoAge acceleration among women, but not among men (women: β = 1.4; 95% CI: −0.4, 3.3 vs. men: β = −0.3; 95% CI: −2.2, 1.5) in fully adjusted models. In models stratified on social cohesion, association of neighborhood poverty and quality with accelerated DNAm aging remained elevated for residents living in neighborhoods with lower social cohesion, but were null for those living in neighborhoods with higher social cohesion. Our study suggests that living in adverse neighborhood conditions can speed up epigenetic aging, while positive neighborhood characteristics may buffer effects.

## INTRODUCTION

Inequalities in aging-related health and mortality based on neighborhood conditions have been consistently shown in the literature, independent of individual-level socioeconomic position (1–8). This has significant implications for Black individuals in the United States who are overrepresented in deprived and socioeconomically disadvantaged neighborhoods due to historical racialized segregation policies and limited upward residential mobility (9–11). Despite the robust evidence linking the neighborhood environment to health and mortality, less is known about the underlying biological mechanisms explaining how neighborhood conditions become physiologically embodied to influence longevity.

Growing evidence suggests that accelerated biological aging at the cellular- and molecular-levels may help to explain the mechanisms linking adverse neighborhood environments to mortality (12–19). Epigenetic mechanisms, particularly DNA methylation (DNAm), has offered promising insights into possible molecular pathways. Specifically, DNAm regulates gene expression without altering the underlying DNA sequence, is associated with health outcomes and mortality, and responsive to exogenous factors, including area-level environmental exposures (15,16,20–22). Several measures of DNAm aging, also known as “epigenetic clocks,” have been developed using DNA microarray technology and statistical algorithms to identify alterations in DNAm that are highly correlated with chronological age (23–25). Recently, Levine et al., developed a DNAm aging biomarker to better estimate aging as related to clinical phenotypes of chronic disease (25). Accelerated DNAm aging, defined as instances where DNAm age is greater than chronological age, has been associated with adverse physical and mental health outcomes and all-cause mortality (26). Taken together, it is possible that molecular biomarkers, such as DNAm, are useful early indicators of neighborhood quality-related health and aging effects.

Living in areas characterized by exposure to disadvantage (18,22,27), violence and crime (28), and air pollution (29) have been shown to accelerate DNAm aging. For studies of neighborhood environment and DNAm aging, neighborhood environment is most often assessed using administrative data sources (i.e. U.S. Census Bureau’s American Community Survey) and boundaries (i.e. census tract). However, an individual’s perception of their neighborhood environment may also capture salient experiences of the neighborhood environment that impact molecular mechanisms (16). Positive aspects of the neighborhood, such as perception of neighborhood social cohesion, may act to buffer the effects of an adverse neighborhood environment on DNAm aging, as found in studies relating neighborhoods to health (30–33). Accordingly, our study sought to investigate both the independent and joint impacts of neighborhood quality and social cohesion on three measures of DNAm aging—Horvath’s epigenetic clock, Hannum’s epigenetic clock, and Levine’s PhenoAge— among a sample of predominately Black adults living in Detroit, MI (23–25). Given previous work suggesting biological aging biomarkers may differ by gender and with respect to area-level environmental characteristics (12,29,34), we investigated associations stratified by gender as well as in the full sample.

## METHODS

### Study population

Baseline data from adults participating in the Detroit Neighborhood Health Study (DNHS) were used for the present study. The DNHS is a population-based prospective cohort study of a representative sample of primarily Black adult residents (18 years of age or older) living in Detroit, MI. Study participants completed annual structured telephone interviews from 2008 through 2012 to assess perceptions of neighborhoods, mental and physical health status, social support, and alcohol and tobacco use. Biospecimens were also collected annually through 2013. A total of 2,081 adults participated in the DNHS, of which 612 were asked to provide either a venipuncture or blood spot at baseline. DNA methylation was measured from whole blood in 179 participants as part of a pilot study, of which 158 with both DNA methylation and neighborhood quality data at baseline examination were included this analysis. Informed consent was received from all participants in the study prior to participation and the DNHS was approved by the institutional review boards at the University of Michigan (HUM00014138) and the University of North Carolina at Chapel Hill (13-3999).

### Neighborhood Quality Assessment

Neighborhood quality was assessed using information from the U.S. American Community Survey (ACS) participant survey responses, objective neighborhood evaluations, and participant’s survey responses. To estimate neighborhood poverty and unemployment, we used data from the 2008-2013 ACS. Neighborhood poverty was defined by the percent of individuals below the federal poverty level. Neighborhood estimates at the census block-group level were aggregated to derive measures within the 54 historically defined neighborhoods in Detroit (35). Neighborhood poverty was converted to z-scores for analyses.

Objective neighborhood measures, conducted by trained personnel, were captured from structured assessments of Detroit’s 54 neighborhoods. Within the 54 neighborhoods, 135 block groups were assessed between June and July 2018. During this time, 19 neighborhood characteristics (**Supplemental Table 1**) were evaluated using a standardized instrument adapted from the New York Social Environment Study to be relevant to Detroit (36). Evaluators responded with “yes” or “no” and the frequency of “yes” responses were calculated for each variable by block group, which were then averaged by neighborhood. As previously described, a principal components (PCs) analysis was performed to summarize the correlated neighborhood characteristics and minimize the number of statistical tests performed due to our limited sample size (18). Based on the screeplot and an *a priori* cutoff of 90% variance explained, eight PCs were selected to assess objective neighborhood quality (**Supplemental Table 2**) (18).

During the first telephone interview, participants were asked to report on their perceptions of their neighborhood community. Participants were asked about the following characteristics: 1) a close-knit or unified neighborhood; 2) people around here are willing to help their neighbors; 3) people in this neighborhood generally don’t get along with each other (reverse scored); 4) people in this neighborhood do not share the same values (reverse scored); 5) people in this neighborhood can be trusted. Each response was coded from *strongly disagree* (0) to *strongly agree* (3). Responses were summed to create a score of individual perception of neighborhood social cohesion, which ranged from 0 to 15 with higher scores indicating greater perception of neighborhood social cohesion. To generate a measure of neighborhood-level perception of social cohesion, individual scores were aggregated and mean neighborhood social cohesion was assigned for each participant living within the same neighborhood (37,38).

### DNAm age variables

Peripheral blood DNA was extracted using venipuncture and genome-wide DNA methylation was measured in whole blood leukocytes using the Illumina Infinium HumanMethylation 450k array using published methods (39). Samples were bisulfite converted using the EZ-96 DNA methylation kit (Zymo Research). Sample quality control protocol excluded samples with probe detection call rates <90% and those with an average intensity value of either <50% or sample mean <2,000 arbitrary units (AU). Quality control was performed using the R package CpGassoc (40). Probes with detection p-values > 0.001 and samples with missing data for > 10% of probes were also removed, along with known SNPs and cross-hybridizing probes (41). Probe normalization was performed using the beta-mixture quantile normalization method using the R package wateRmelon. (42,43). Following normalization, ComBat was used to account for batch effects using *M* values converted from the beta values (44). *M* values were converted back to beta values for calculation of DNAm age variables.

Horvath’s epigenetic clock, Hannum’s epigenetic clock, and Levine’s PhenoAge were calculated using their published algorithms (23–25). Horvath’s epigenetic clock consists of 353 CpG sites and is applicable across multiple sources of cells, tissues, and organs across the entire lifespan, including whole blood in adults (24). Hannum’s epigenetic clock is a single tissue estimator derived from 71 CpG sites from DNA of adult whole blood samples (23). Horvath’s and Hannum’s clocks are both highly correlated with chronological age; however, weaker associations are observed with clinical characteristics of physiological dysfunction (26). Levine’s PhenoAge measure was developed using 513 epigenetic loci associated with 10 clinical phenotypes, – albumin, creatinine, glucose, C-reactive protein, lymphocyte percent, red blood cell volume, red cell distribution, alkaline phosphatase, white blood cell count, and chronological age – which were further validated for associations with mortality, co-morbid disease burden, and physical function (25). The 513 epigenetic loci were converted into an epigenetic clock using regression modeling as done for previous clocks. Levine’s PhenoAge estimator (and PhenoAge acceleration) strongly predicts several aging-related outcomes including all-cause mortality, cancer, coronary heart disease, and Alzheimer’s disease. This resulted in the selection of 513 CpG sites for estimation of phenotypic age. Each measure of DNAm age was strongly correlated with chronological age in our study population (**Supplemental Figures 1A-C**). For the purpose of this analysis, DNAm age residuals were calculated by regressing each DNAm age variable on chronological age resulting in positive and negative deviations from chronological years of age. Positive scores (residuals from the regression model) reflected accelerated DNAm aging.

### Additional Covariates

Baseline demographic, social, and behavioral factors associated with neighborhood quality and DNAm aging were collected, including chronological age, race/ethnicity, educational attainment, employment status, and number of years lived in current neighborhood. Chronological age was self-reported and included in models as a continuous variable. Race/ethnicity, educational attainment, and employment status were also self-reported and were categorized as shown in **Table 1**. Participants reported the number of years lived in current neighborhood, which was treated as a continuous variable. Current smoking status (never, ever, current) and lifetime alcohol intake (ever vs. never) were also included in the models.

**Table 1.**
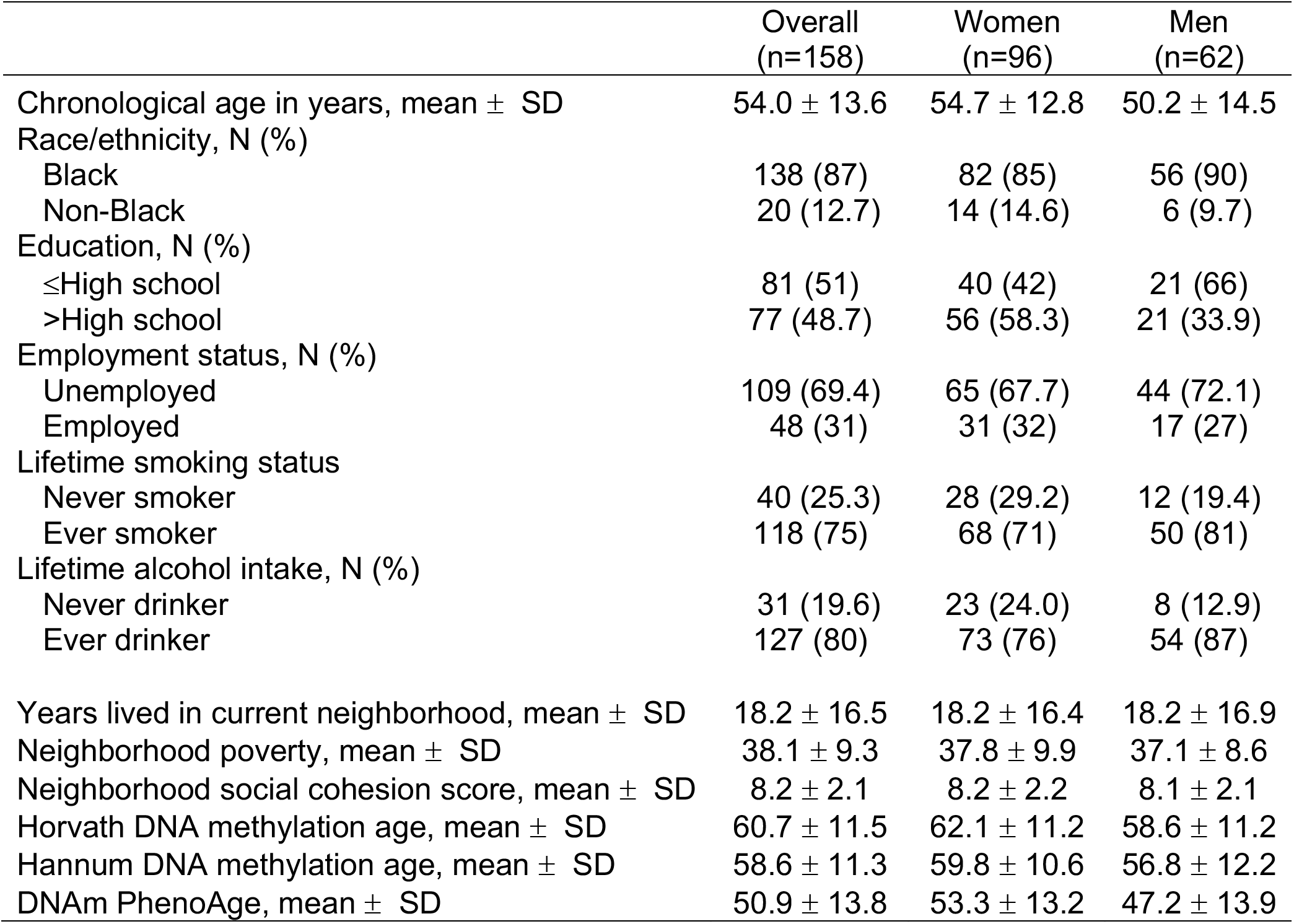
Selected sociodemographic and lifestyle characteristics of 158 participants in the Detroit Neighborhood Health Study

### Statistical Analysis

Distributions of select baseline covariates were examined. Three gender-specific age acceleration variables – Horvath age acceleration (from Horvath’s clock), Hannum age acceleration (from Hannum’s clock), and PhenoAge acceleration (from Levine’s clock) – were derived from regression residuals for the full sample and separately for men and women. We examined whether each neighborhood quality measure – neighborhood poverty, objective neighborhood evaluation (PCs), and neighborhood social cohesion — were independently associated with accelerated DNAm aging. Two separate models were used to estimate our associations of interest: (1) the unadjusted model of neighborhood quality and DNAm age acceleration; and (2) the full model adjusted for race/ethnicity, educational attainment, employment status, lifetime smoking, lifetime alcohol intake, and number of years residing in current neighborhood. We also examined these associations stratified by gender. In a second set of analyses, we assessed whether perception of neighborhood social cohesion provided a buffering effect of the neighborhood poverty and quality measures on DNAm aging. To estimate this, full sample models were stratified by a binary indicator of neighborhood social cohesion, categorized as higher vs. lower social cohesion using the median value. We did not examine this association in the gender-specific models due to our limited sample sizes. All analyses were performed in R version 3.6 (45). Results are reported using regression coefficient (ß) and associated 95% confidence interval (CI). Given the small sample size and correlations, as well as biological relations, among the epigenetic aging outcome measures we did not impose a multiple testing correction.

## RESULTS

### Description of study sample

Chronological age was highly correlated with Horvath’s clock (Pearson r=0.81), Hannum’s clock (Pearson r=0.84), and Levine’s clock (Pearson r=0.80; **Supplemental Figure 1**). Selected baseline characteristics for the 158 study participants included in this analysis are shown in **Table 1**. Overall, the study sample comprised primarily of Black (87%) and women (61%) participants. Participants were long-term residents of their neighborhoods (mean = 18.2 ± 16.5 years). The mean proportion of residents living below the federal poverty level was 38.1% ± 9.3%, while the mean social cohesion score was 8.2 (SD = 2.1). On average, men were 4.5 years younger than women (men: 50.2 years ± 14.5; women: 54.7 years ± 12.8). For both men and women, DNAm age was higher for Horvath’s clock (men: 58.6 ± 11.2 years; women: 62.1 ± 11.2 years) and Hannum’s clock (men: 56.8 ± 12.2; women: 59.8 ± 10.6 years) and lower for Levine’s clock (men: 47.2 ± 13.9; women: 53.3 ± 13.2 years).

### Neighborhood poverty and DNAm age

We observed evidence to indicate neighborhood poverty was associated with DNAm age acceleration that was primarily driven by the association among women (**Figure 1**). After adjusting for factors included in the full model, the pattern of association among women signaled living in neighborhoods with higher poverty was associated with PhenoAge acceleration, which was not seen among men (women: β = 1.4; 95% CI: −0.4, 3.3 vs. men: β = −0.3; 95% CI: −2.2, 1.5). Similar association patterns were observed between neighborhood poverty and Horvath and Hannum age acceleration among women. The associations among men were primarily null (**Supplemental Table 3**).

**Figure 1.**
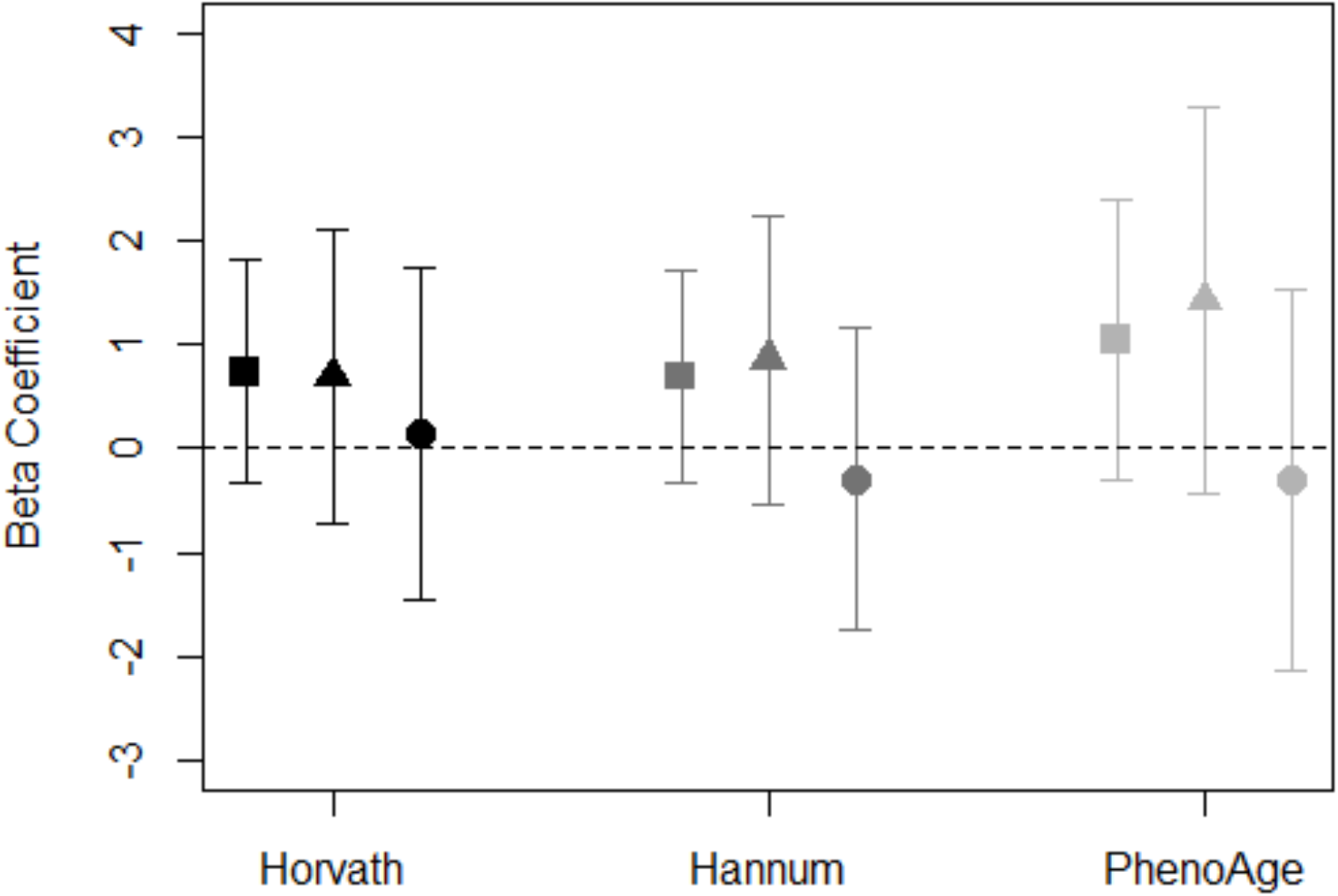
Association between neighborhood poverty and DNAm age acceleration measures for total sample (square), women (triangle), and men (circle). Models adjusted for race/ethnicity, education level, employment, smoking status, alcohol intake, and years residing in current neighborhood. Black symbols represent associations with Horvath age acceleration, dark gray represent Hannum age acceleration, and light gray represent PhenoAge acceleration.

### Objective neighborhood observations (PCs) and DNAm age

Of the eight principal components (PCs) retained, only PC7 showed a consistent pattern of association with accelerated aging (**Supplemental Table 4**). Among the total sample, PC7 was associated with accelerated DNAm aging for Horvath age acceleration (β = 1.8; 95% CI: 0.4, 3.1), Hannum age acceleration (β = 1.7; 95% CI: 0.4, 3.0), and PhenoAge acceleration (β = 2.1; 95% CI: 0.4, 3.8) in the full model (**Figure 2**). The top positive loadings for PC7 were factors characterized by the presence of abandoned cars and people on the street (**Supplemental Table 2**). Among women, we observed PhenoAge acceleration in response to living in a neighborhood characterized by PC7 (β = 2.4; 95% CI: −0.0, 4.9), and, among men, suggestion of Horvath age acceleration (β = 1.9; 95% CI: −0.1, 4.0; **Figure 2, Supplemental Table 4**).

**Figure 2.**
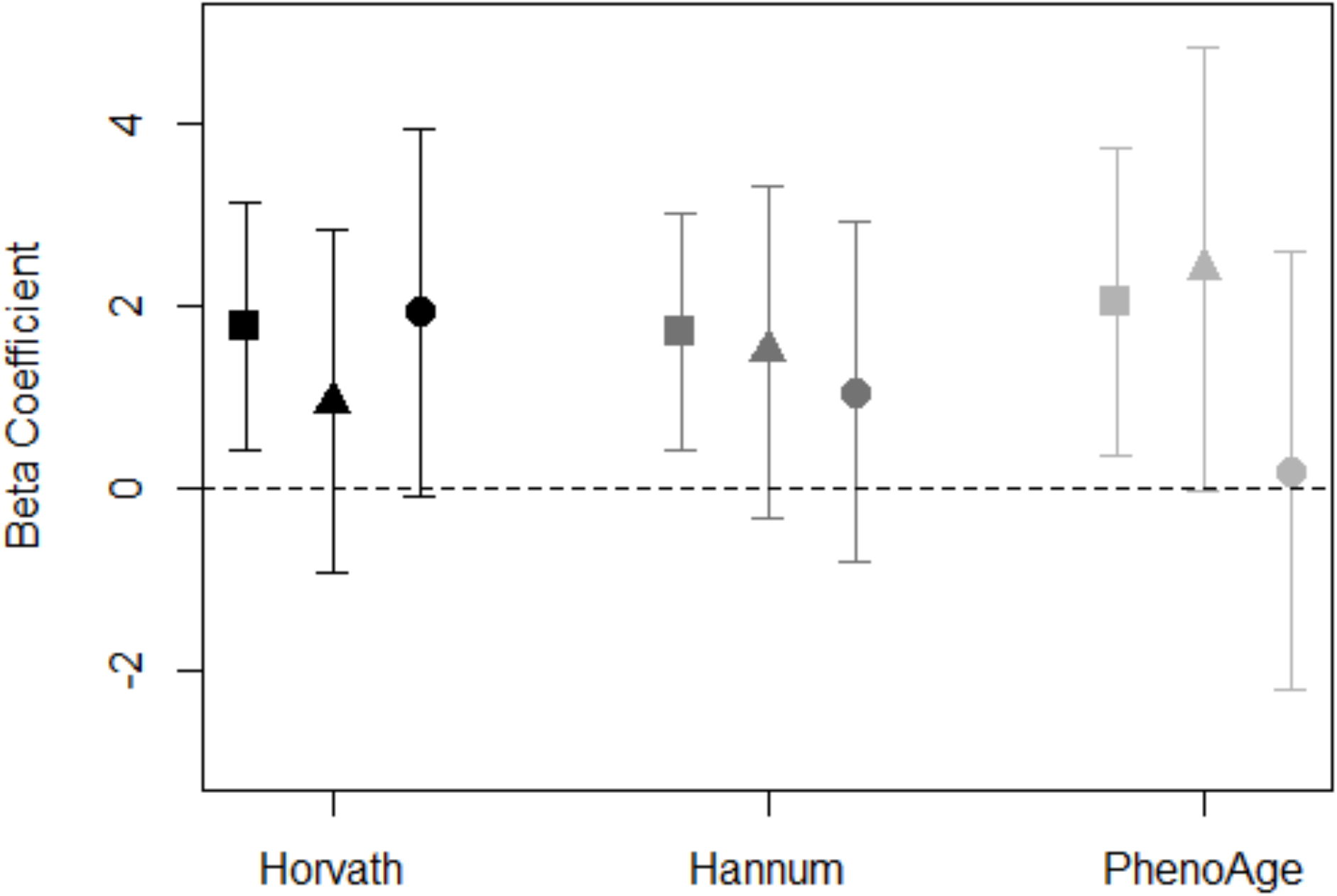
Association between neighborhood PC7 and DNAm age acceleration measures for total sample (square), women (triangle), and men (circle). Models adjusted for race/ethnicity, education level, employment, smoking status, alcohol intake, and years residing in current neighborhood. Black symbols represent associations with Horvath age acceleration, dark gray represent Hannum age acceleration, and light gray represent PhenoAge acceleration.

### Neighborhood social cohesion and DNAm age

The associations in the full sample between neighborhood social cohesion and DNAm aging biomarkers were largely null (**Figure 3**). However, we observed potential differences by gender. For instance, among men, neighborhood social cohesion was associated with negative age acceleration for Hannum age acceleration (β = −0.6; 95% CI: −1.2, 0.1) and PhenoAge acceleration (β = −0.7; 95% CI: −1.6, 0.1); whereas, the effect estimates for associations between neighborhood social cohesion with Hannum age acceleration and PhenoAge acceleration were null for women (**Supplemental Table 3**).

**Figure 3.**
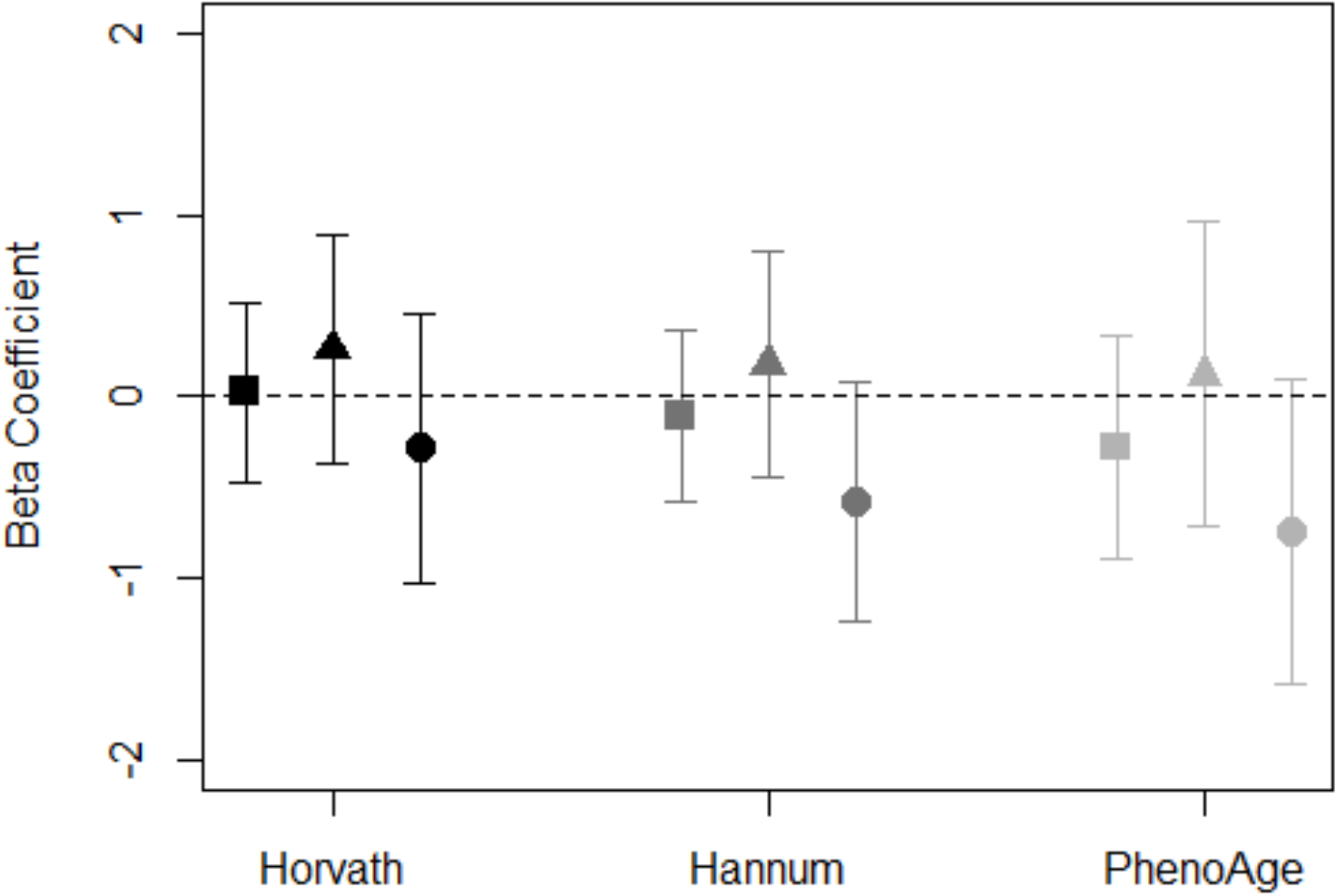
Association between neighborhood social cohesion and DNAm age acceleration measures for total sample (square), women (triangle), and men (circle). Models adjusted for race/ethnicity, education level, employment, smoking status, alcohol intake, and years residing in current neighborhood. Black symbols represent associations with Horvath age acceleration, dark gray represent Hannum age acceleration, and light gray represent PhenoAge acceleration.

To assess whether objective neighborhood quality-DNAm aging associations differed by neighborhood social cohesion in the full sample, we conducted stratified analyses in the full models comparing associations for individuals living in neighborhoods with higher vs. lower social cohesion scores (based on the median value of 8.1). Our results were suggestive of a difference in the neighborhood PC7-DNAm age acceleration association by neighborhood social (**Figure 4**). We found that the association between PC7 and DNAm age acceleration remained among participants living in neighborhoods with lower social cohesion elevated (Horvath: β = 2.1; 95%. CI: 0.6, 3.6; Hannum: β = 2.2; 95%. CI: 0.6, 3.8; PhenoAge: β = 2.3; 95%. CI: 0.3, 4.7); however, there appeared to be no association among participants living in neighborhoods with higher neighborhood social cohesion (**Figure 4, Supplemental Table 5**). Similar patterns of association were observed for neighborhood poverty and DNAm age acceleration for participants living in neighborhoods with lower social cohesion; however, null associations were observed among residents living in neighborhoods with higher social cohesion (**Supplemental Figure 2**).

**Figure 4.**
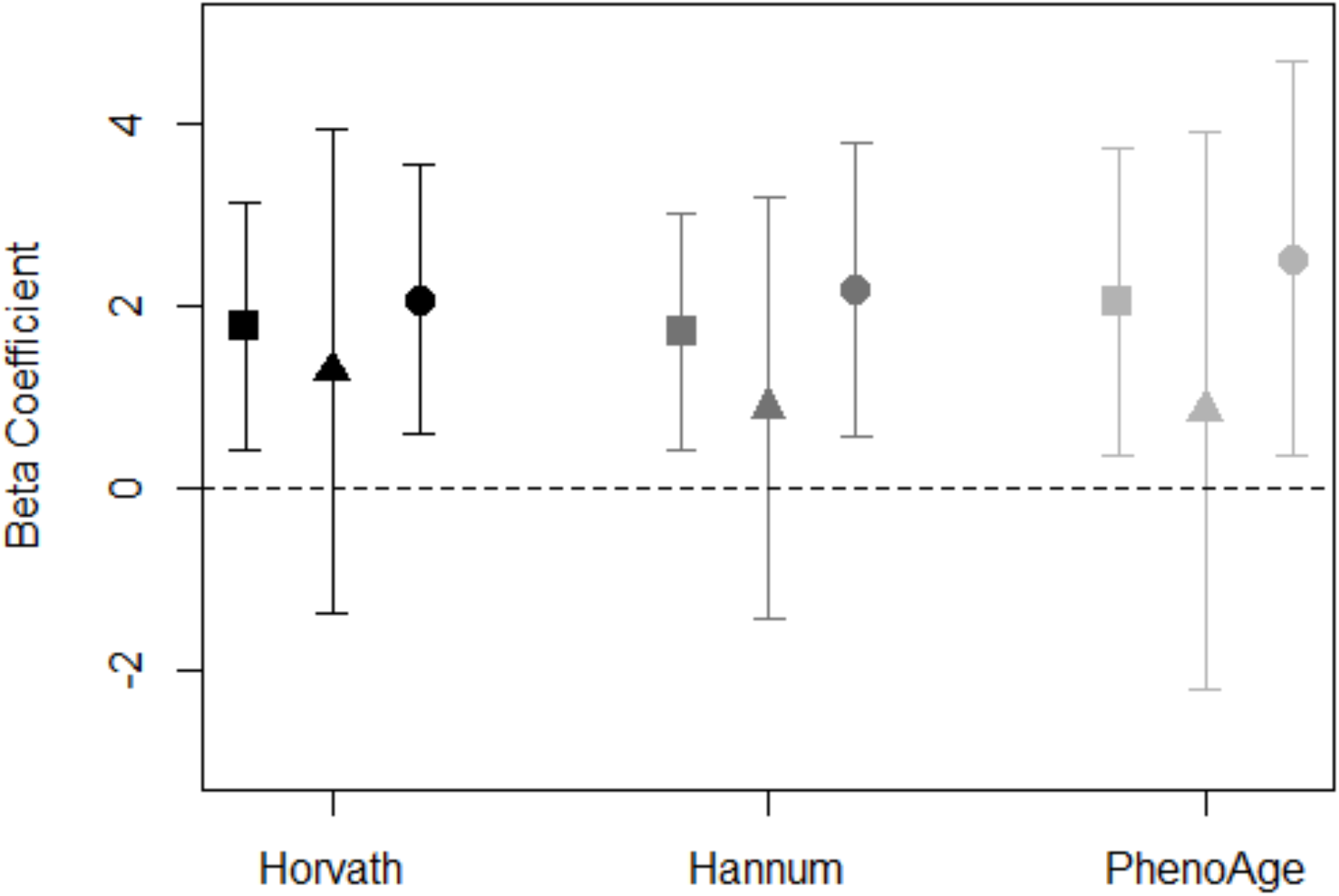
Association of PC7 and DNAm age acceleration measures stratified by neighborhood social cohesion for total sample (square), high social cohesion (triangle), and low social cohesion (circle). Models adjusted for race/ethnicity, education level, employment, smoking status, alcohol intake, and years residing in current neighborhood. Black symbols represent associations with Horvath age acceleration, dark gray represent Hannum age acceleration, and light gray represent PhenoAge acceleration.

## DISCUSSION

In this study of predominately Black adults living in Detroit, we observed associations between neighborhood poverty and aspects of neighborhood quality with accelerated DNAm aging that were primarily driven by associations in women. PhenoAge outpaced chronological age by nearly 2 years for women living in neighborhoods with higher measures of poverty and those characterized by increased presence of abandoned cars and people on the street. Neighborhood social cohesion was associated with negative DNAm age acceleration (or DNAm age deceleration) among men, but not women. Further, in models stratified on perceptions of neighborhood social cohesion, associations between objective measures of neighborhood quality—poverty and PC7— and DNAm age acceleration remained elevated for participants living in neighborhoods with lower social cohesion, while associations were relatively null for those residing in neighborhoods with high social cohesion. Taken together, it is possible that living in neighborhoods with higher social cohesion, despite higher poverty and disorder, may protect against accelerating DNAm aging.

Our findings are consistent with previous research linking neighborhood disadvantage to accelerated DNAm aging in children and adults (22,27). In our study, we found that women’s DNAm aging was more sensitive to neighborhood poverty and PC7 than men. Similarly, a prior study among 100 African American women found that a standard deviation increase in neighborhood disadvantage was associated with a 9-month increase (β = 0.75; 95% CI: 0.27, 1.23) in Hannum’s age acceleration (22), which is comparable to the 12-month increase in Hannum’s age acceleration we observed in relation to a standard deviation increase in neighborhood poverty among women in our study. Additionally, we explored the impact of neighborhood poverty on a more recent measure of epigenetic aging, Levine’s clock, which estimates aging based on epigenetic loci associated with 10 age-related clinical measures (25). Neighborhood poverty appears to have a greater effect on PhenoAge acceleration than Hannum’s age acceleration in our study. Many of the clinical measures included in Levine’s PhenoAge estimator, such as glucose levels (46) and C-reactive protein (47–49), have been shown to be sensitive to neighborhood-level social characteristics, which might make PhenoAge more sensitive to the social characteristics and explain stronger associations with PhenoAge than observed for other epigenetic aging biomarkers in this and previous studies of the neighborhood social environment.

Unlike previous studies, our study used direct observations of neighborhood aspects conducted by trained evaluators, which provide an objective assessment of the neighborhood’s natural social environment (50). To reduce the number of tests conducted on our limited sample size, we summarized the 19 observed indicators of neighborhood quality using PCA and found that PC7, defined by the presence of people on the street and abandoned cars, was associated with accelerated DNAm aging across each of the three measures of epigenetic aging examined in this study. This finding is consistent with our previous work showing neighborhood quality measures, namely abandoned cars, people on the street, and non-art graffiti, were associated with an epigenetic biomarker of mortality (18).

Neighborhood social cohesion has also been shown to affect physical and mental health outcomes and may have effects at the molecular levels (16,30–33). In a study of 1,226 adults in the Multi-Ethnic Study of Atherosclerosis (MESA), the neighborhood social environment, measured as a summary score that included social cohesion, was associated with DNAm in stress- and inflammation-related genes (16). In our study, men living in neighborhoods with higher neighborhood social cohesion appeared to experience negative Hannum and Pheno age acceleration. Furthermore, in our full models stratified by higher vs. lower neighborhood social cohesion, the impact of PC7 and poverty on DNAm aging remained detrimental among participants living in neighborhoods with lower social cohesion, but not among those living in neighborhoods with higher social cohesion. Additional studies in larger prospective cohorts are needed to corroborate our findings; yet, if replicated our results have important implications for mortality and longevity. Evidence exists demonstrating that neighborhood social cohesion is associated with lower risk of mortality (1). Given accelerated DNAm aging’s link to mortality, it is possible residing in a neighborhood with greater social cohesion could buffer the negative effects of neighborhood deprivation on mortality through decelerated DNAm aging as demonstrated in our study.

Our study examines the impact of neighborhood level social processes on individual-level molecular mechanisms. Several potential mechanisms exist that may explain molecular response to the neighborhood environment. First, participants residing in neighborhoods with high levels of disadvantage and disorder may experience chronic activation and dysregulation of stress-response and inflammatory pathways and altered DNAm patterns, as suggested in previous studies demonstrating associations between neighborhood disadvantage and DNAm of genes involved in stress-response and inflammation (15,16). Additional pathways, such as smoking status and dietary behaviors, may affect these associations. Unfortunately, we did not have information on dietary habits, but did adjust for lifetime smoking and alcohol consumption in our models, which did not impact our results. There may be additional factors correlated with neighborhood disadvantage like environmental air quality that may explain our associations and should be explored in future investigations. We also observed distinct patterns of associations for men and women. While these findings warrant follow-up in larger samples, the null associations observed between neighborhood poverty and PC7 and DNAm aging acceleration may be explained by the overall health of men included in our small study sample. For instance, PhenoAge – a marker of the aging process for important clinical measures of chronic disease – was lower among men in our study than women, which may be a signal that men were in better overall health, and DNAm aging was not responsive to the adverse effects of living in neighborhoods with poor economic and social conditions. Future studies should examine this further.

Our study is not without limitations. Due to our limited sample size these analyses should be considered somewhat exploratory in nature. We did not impose a multiple testing correction as the sample size was limited and the outcomes were correlated, as well as biologically related. In addition, due to the limited sample size, we were unable to explore potentially salient higher-order interactions, such as three-way interactions between gender, neighborhood disadvantage, and social cohesion, which are likely relevant but cannot be addressed using small sample sizes. Despite our sample size, we were able to pick up signals of associations in the entire cohort and when stratifying on gender and neighborhood social cohesion. We focused on summary measures of objective neighborhood quality as these summary measures may combine correlated features of the underlying inputs which can increase power to detect associations. DNHS is a population-based study of individuals residing in the Detroit metropolitan area, which may limit the generalizability of our study results to other parts of the United States. However, our findings are in line with previous studies of neighborhood disadvantage and DNAm aging. Our analysis is cross-sectional with neighborhood measures and DNAm assessed at similar points in time restricting us from being able to establish temporal sequence. However, it is unlikely that individuals selected into their neighborhoods based on their underlying epigenetic age, particularly given that most individuals resided in their homes for over a decade. When combined with the average age of individuals (54.0), this would indicate that most individuals began residing in their homes at an age prior to when age (or epigenetic age) driven functional deficits would appear, which would have been the most likely causal mechanism for epigenetic age acceleration to drive neighborhood choice. Detroit was one of the harder hit areas during to the U.S. Great Recession from 2007 to 2009, which coincided with the DNHS. While participants in the neighborhoods were long-term residents, we do not have repeated measures of neighborhood quality and are unable to examine the effects of neighborhood changes on DNAm.

In summary, our study explored multiple measures of neighborhood environment, including U.S. Census data for neighborhood poverty, direct observations of neighborhood quality, and survey responses of neighborhood social cohesion, in relation to DNAm aging. We found that individuals living in neighborhoods characterized by higher levels of poverty and disorder may experience accelerated aging, where their DNAm age is higher than their chronological age. Yet, effects of neighborhood poverty and quality on DNAm aging may be buffered by increased neighborhood social cohesion. Given that communities of color are more likely to reside in deprived and disadvantaged neighborhoods, our findings offer a molecular insight into potential mechanisms of health disparities. We suggest that future studies fruitfully interrogate these associations to build upon our understanding of the biosocial mechanisms that contribute to racial/ethnic disparities in health.

## ACKNOWLEDGEMENTS

We are grateful for the time and effort of the DNHS study participants, team, and volunteers. The study authors would also like to acknowledge Sandra Alvarez and Cierra Dungee for their preliminary work on epigenetic aging and neighborhood principal components. This work was supported by multiple grants from the National Institutes of Health (R01D022720, R01DA022720-S1, RC1MH088283, R01MD011728). CLM received salary and training support from Eunice Kennedy Shriver National Institute of Child Health and Human Development grant T32HD007168, K99MD012808, and R01MD013349. TMZ received support from an NSF Career Award (grant no. 1845796). This manuscript does not necessarily represent the views and policies of the US Environmental Protection Agency. Any mention of tradenames or copyright does not constitute endorsement by the US Environmental Protection Agency.

## Notes

### Competing Interest Statement

The authors have declared no competing interest.

