## Supplemental Figures for "Neighborhood environment, social cohesion, and epigenetic aging"

**Supplemental Figure 1.** Correlations between chronological age and A) Horvath's DNAm age, B) Hannum's DNAm age, and C) Levine's clock.

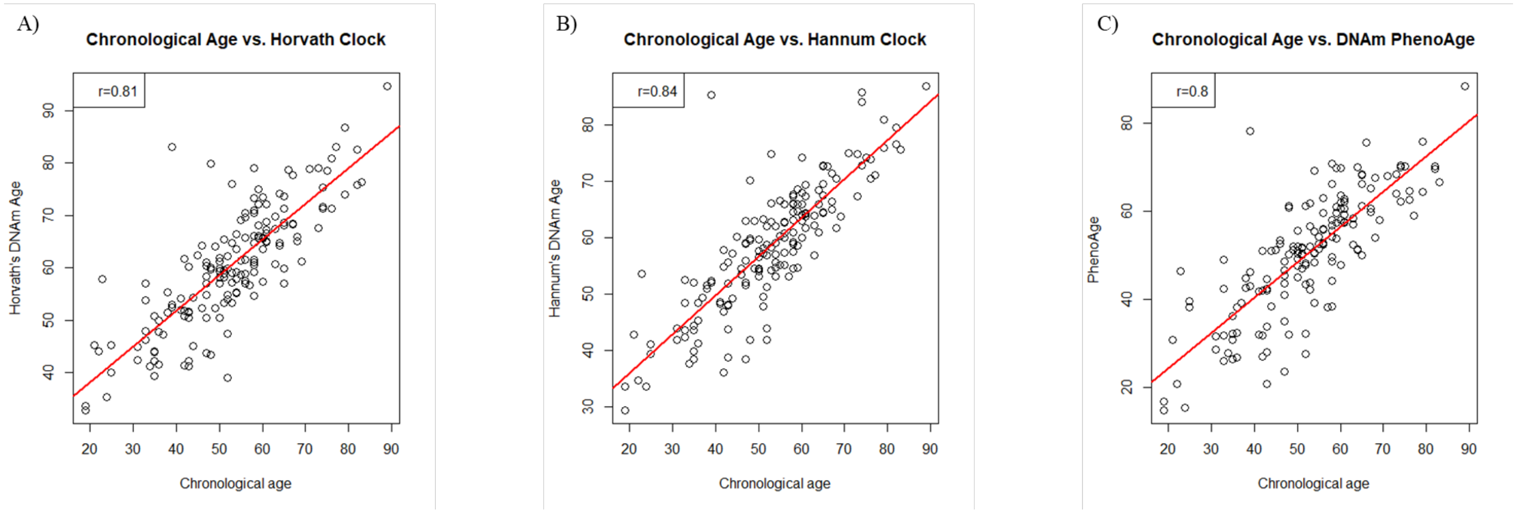

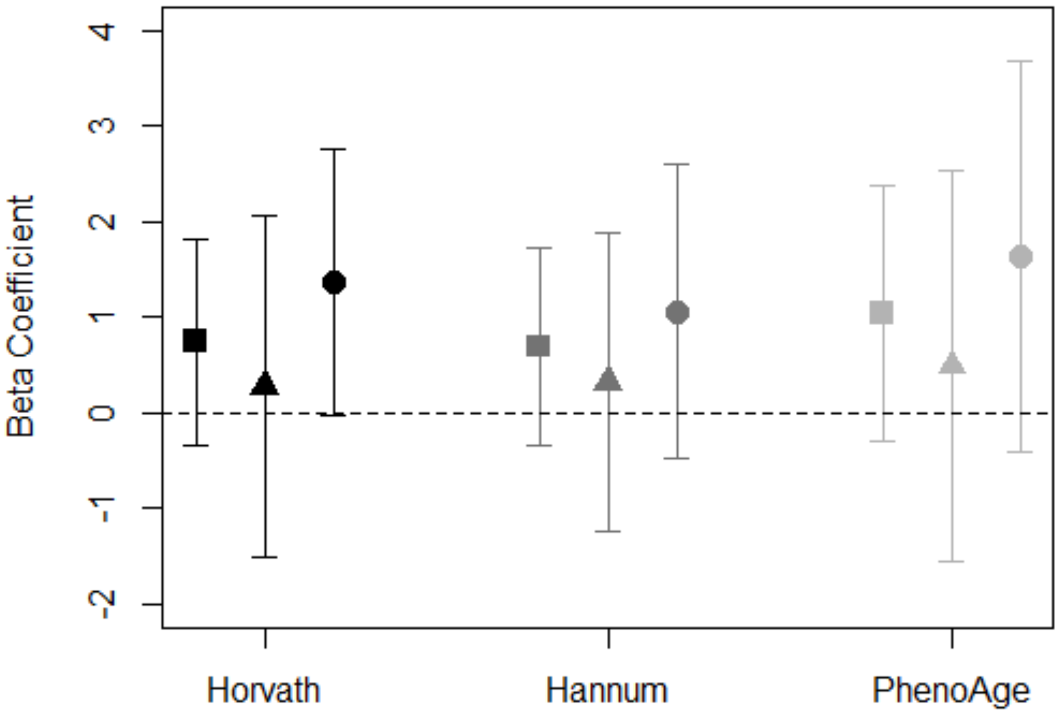


**Supplemental Figure 2.** Association between neighborhood poverty and DNAm aging acceleration measures stratified by neighborhood social cohesion for total sample (square), high social cohesion (triangle), and low social cohesion (circle). Models adjusted for race/ethnicity, education level, employment, smoking status, alcohol intake, and years residing in current neighborhood. Black symbols represent associations with Horvath age acceleration, dark gray represent Hannum age acceleration, and light gray represent PhenoAge acceleration
