## Supplemental Tables for "Neighborhood environment, social cohesion, and epigenetic aging"

**Supplemental Table 1**. Definitions and Distributions (mean and standard deviation (SD)) for the 19 neighborhood quality indicators assessed by two trained assessors for a select number of census block groups in the Detroit Neighborhood Health Study.

| Neighborhood Quality Indicator | Metric as Evaluated by Trained Assessor | Mean | SD |
|---|---|---|---|
| HQ1 | Are there any buildings with broken windows, boarded up windows, or boarded up doors? Percent of sampled block group segments within Neighborhood that have "Yes" for this question. | 34.2 | 14.1 |
| HQ2 | Are there any buildings with outside damage that can only be corrected by major repairs such as siding, shingles, boards, brick, concrete, and stucco? Percent of sampled block group segments within Neighborhood that have "Yes" for this question. | 29.7 | 14 |
| HQ3 | Are there any entirely vacant buildings? Percent of sampled block group segments within Neighborhood that have "Yes" for this question. | 34 | 11.8 |
| HQ4 | Are there any empty, vacant lots? Percent of sampled block group segments within Neighborhood that have "Yes" for this question. | 32.6 | 21.4 |
| HQ5 | Are there any construction sites? Percent of sampled block group segments within Neighborhood that have "Yes" for this question. | 2.4 | 2.5 |
| HQ6 | Is there a community garden? Percent of sampled block group segments within Neighborhood that have "Yes" for this question. | 0.566 | 0.873 |
| HQ7 | Is there graffiti (non-art)? Percent of sampled block group segments within Neighborhood that have "Yes" for this question. | 17.2 | 10.3 |
| HQ8 | Are the street and sidewalk clean? Percent of sampled block group segments within Neighborhood that have "Yes" for this question. | 73.3 | 14.1 |
| HQ9 | Are there any big, mature trees? Percent of sampled block group segments within Neighborhood that have "Yes" for this question. | 83.1 | 9.64 |

| | | | |
|---|---|---|---|
| HQ10 | Is there heavy traffic volume? Percent of sampled block group segments within Neighborhood that have "Yes" for this question. | 30.2 | 10.8 |
| HQ11 | Is the street in poor condition? Percent of sampled block group segments within Neighborhood that have "Yes" for this question. | 33 | 11.4 |
| HQ12 | Is the sidewalk in poor condition? Percent of sampled block group segments within Neighborhood that have "Yes" for this question. | 58.7 | 15.2 |
| HQ13 | Is the street noisy? Percent of sampled block group segments within Neighborhood that have "Yes" for this question. | 26.9 | 10.3 |
| HQ14 | Are there people visible on the street? Percent of sampled block group segments within Neighborhood that have "Yes" for this question. | 13.7 | 8.68 |
| HQ15 | Are there any abandoned cars? Percent of sampled block group segments within Neighborhood that have "Yes" for this question. | 6.57 | 4.04 |
| HQ16 | Are any of the following signs visible? A. Neighborhood or Crime Watch. B. Security warning signs. Percent of sampled block group segments within Neighborhood that have "Yes" for this question. | 51.8 | 14.4 |
| HQ17 | Are there any tobacco product advertising signs visible? Percent of sampled block group segments within Neighborhood that have "Yes" for this question. | 1.65 | 1.88 |
| HQ18 | Are there any alcohol advertising signs visible? Percent of sampled block group segments within Neighborhood that have "Yes" for this question. | 2.49 | 2.01 |
| HQ19 | Are there any For Sale OR For Lease OR For Rent signs visible? Percent of sampled block group segments within Neighborhood that have "Yes" for this question. | 18.6 | 6.72 |

**Supplemental Table 2**. Loadings for each of the top seven principal components.

| | PC1 | PC2 | PC3 | PC4 | PC5 | PC6 | PC7 | PC8 |
|---|---|---|---|---|---|---|---|---|
| HQ1 | -0.35 | 0.10 | -0.08 | 0.06 | -0.12 | 0.07 | -0.02 | 0.00 |
| HQ2 | -0.36 | 0.10 | 0.00 | -0.03 | -0.11 | 0.05 | 0.04 | 0.21 |
| HQ3 | -0.29 | 0.19 | -0.18 | 0.08 | -0.15 | 0.20 | 0.00 | -0.14 |
| HQ4 | -0.35 | -0.04 | 0.08 | -0.14 | 0.17 | 0.07 | -0.26 | 0.15 |
| HQ5 | -0.02 | -0.16 | -0.39 | 0.16 | 0.43 | 0.48 | 0.10 | -0.08 |
| HQ6 | -0.08 | -0.01 | 0.14 | 0.73 | 0.18 | -0.25 | 0.12 | -0.30 |
| HQ7 | -0.23 | -0.21 | 0.19 | 0.34 | -0.23 | 0.15 | 0.08 | 0.35 |
| HQ8 | 0.33 | 0.06 | 0.01 | 0.21 | -0.18 | -0.06 | 0.22 | -0.04 |
| HQ9 | 0.12 | 0.28 | 0.02 | 0.16 | 0.57 | -0.25 | -0.16 | 0.59 |
| HQ10 | 0.01 | -0.45 | 0.11 | -0.10 | 0.27 | 0.07 | 0.11 | -0.02 |
| HQ11 | -0.30 | 0.07 | 0.11 | 0.20 | 0.16 | -0.16 | -0.06 | -0.28 |
| HQ12 | -0.32 | 0.20 | 0.03 | -0.06 | 0.15 | 0.08 | 0.02 | -0.05 |
| HQ13 | 0.03 | -0.47 | 0.02 | -0.02 | 0.20 | 0.14 | -0.11 | -0.20 |
| HQ14 | -0.06 | -0.42 | -0.03 | 0.10 | -0.16 | -0.12 | 0.34 | 0.43 |
| HQ15 | -0.22 | 0.12 | -0.21 | -0.23 | 0.19 | -0.20 | 0.75 | -0.03 |
| HQ16 | 0.28 | 0.24 | -0.22 | 0.01 | 0.03 | 0.05 | 0.14 | -0.03 |
| HQ17 | -0.05 | -0.17 | -0.52 | -0.07 | 0.08 | -0.30 | -0.20 | -0.02 |
| HQ18 | -0.13 | -0.20 | -0.40 | 0.05 | -0.20 | -0.50 | -0.22 | -0.05 |
| HQ19 | 0.08 | 0.09 | -0.45 | 0.33 | -0.16 | 0.35 | -0.08 | 0.20 |

**Supplemental Table 3.** Association between neighborhod characteristics aand epigenetic aging[A]

| | **All** | **Women** | **Men** |
|---|---|---|---|
| | β (95% CI) | β (95% CI) | β (95% CI) |
| | ***Neighborhood Poverty*** | | |
| Horvath age acceleration | 0.7 (-0.3, 1.8) | 0.7 (-0.7, 2.1) | 0.1 (-1.5, 1.7) |
| Hannaum age acceleration | 0.7 (-0.3, 1.7) | 0.9 (-0.5, 2.3) | -0.3 (1.7, 1.2) |
| PhenoAge acceleration | 1.0 (-0.3, 2.4) | 1.4 (-0.4, 3.3) | -0.3 (-2.2, 1.5) |
| | ***Neighborhood Social Cohesion*** | | |
| Horvath age acceleration | 0.0 (-0.5, 0.5) | 0.3 (-0.4, 0.9) | -0.3 (-1.0, 0.5) |
| Hannaum age acceleration | -0.1 (-0.6, 0.4) | 0.2 (-0.5, 0.8) | -0.6 (-1.2, 0.1) |
| PhenoAge acceleration | -0.3 (0.9, 0.3) | 0.1 (-0.7, 1.0) | -0.7 (-1.6, 0.1) |

[A]Models are adjusted for race/ethnicity, education level, employment, smoking status, alcohol intake, and years residing in current neighborhood

**Supplemental Table 4.** Association between principal components and epigenetic aging[A]

| | **All** | **Women** | **Men** |
|---|---|---|---|
| | β (95% CI) | β (95% CI) | β (95% CI) |
| | ***Horvath age acceleration*** | | |
| PC1 | -0.1 (-0.6, 0.3) | -0.1 (-0.6, 0.4) | 0.2 (-0.5, 0.9) |
| PC2 | 0.0 (-0.5, 0.6) | -0.1 (-0.8, 0.6) | 0.1 (-0.7, 0.9) |
| PC3 | 0.7 (-0.1, 1.3) | 0.7 (-0.2, 1.6) | -0.1 (-1.1, 1.0) |
| PC4 | -0.3 (-1.2, 0.7) | 0.0 (-1.3, 1.3) | -0.4 (-1.8, 1.0) |
| PC5 | 0.0 (-1.1, 1.1) | 0.2 (-1.2, 1.6) | 0.2 (-1.2, 1.6) |
| PC6 | 1.0 (-0.2, 2.1) | **1.6 (0.0, 3.1)** | 0.0 (-1.7, 1.8) |
| PC7 | **1.8 (0.4, 3.1)** | 0.9 (-0.9, 2.8) | 1.9 (-0.1, 4.0) |
| PC8 | 0.1 (-1.4, 1.7) | -0.1 (-2.1, 1.8) | -0.6 (-3.0, 1.9) |
| | ***Hannum age acceleration*** | | |
| PC1 | -0.2 (-0.6, 0.2) | -0.2 (-0.7, 0.3) | 0.3 (-0.3, 0.8) |
| PC2 | 0.1 (-0.4, 0.6) | -0.1 (-0.8, 0.6) | 0.3 (-0.5, 1.0) |
| PC3 | 0.3 (-0.3, 1.0) | 0.2 (-0.7, 1.1) | -0.0 (-1.0, 0.9) |
| PC4 | -0.1 (-1.0, 0.7) | 0.0 (-1.2, 1.3) | -0.1 (-1.4, 1.2) |
| PC5 | -0.0 (-1.1, 1.0) | 0.1 (-1.2, 1.5) | 0.8 (-0.8, 2.4) |
| PC6 | 0.4 (-0.8, 1.5) | 1.1 (-0.5, 2.6) | -0.7 (-2.3, 0.9) |
| PC7 | **1.7 (0.4, 3.0)** | 1.5 (-0.3, 3.3) | 1.1 (-0.8, 2.9) |
| PC8 | 0.3 (-1.2, 1.8) | 0.9 (-1.0, 2.8) | -1.9 (-4.1, 0.2) |
| | ***PhenoAge acceleration*** | | |
| PC1 | -0.3 (-0.8, 0.2) | -0.3 (-1.0, 0.4) | 0.3 (-0.5, 1.0) |
| PC2 | -0.1 (-0.8, 0.6) | -0.4 (-1.3, 0.5) | 0.1 (-0.8, 1.0) |
| PC3 | 0.4 (-0.4, 1.3) | 0.3 (-0.9, 1.5) | -0.1 (-1.8, 0.5) |
| PC4 | -0.6 (-1.8, 0.5) | -0.2 (-1.8, 1.5) | -0.9 (-2.5, 0.7) |
| PC5 | -0.6 (-1.9, 0.7) | -0.6 (-2.4, 1.2) | -0.2 (-2.3, 1.9) |
| PC6 | 0.4 (-1.1, 1.9) | 1.5 (-0.6, 3.6) | -1.0 (-3.0, 1.0) |
| PC7 | **2.1 (0.4, 3.8)** | 2.4 (-0.0, 4.9) | 0.2 (-2.2, 2.6) |
| PC8 | -0.0 (-1.9, 1.9) | -0.2 (-2.8, 2.4) | -0.8 (-3.7, 2.0) |

[A]Models are adjusted for race/ethnicity, education level, employment, smoking status, alcohol intake, and years residing in current neighborhood

**Supplemental Table 5.** Association between neighborhod characteristics and epigenetic aging by neighborhood social cohesion[A]

| | **All** | **High Social Cohesion** | **Low Social Cohesion** |
|---|---|---|---|
| | β (95% CI) | β (95% CI) | β (95% CI) |
| ***Neighborhood Poverty*** | | | |
| Horvath age acceleration | 0.7 (-0.3, 1.8) | 0.3 (-1.5, 2.1) | 1.3 (-0.0, 2.8) |
| Hannaum age acceleration | 0.7 (-0.3, 1.7) | 0.3 (-1.2, 1.9) | 1.1 (-0.5, 2.6) |
| PhenoAge acceleration | 1.0 (-0.3, 2.4) | 0.5 (-1.6, 2.5) | 1.6 (-0.4, 3.7) |
| ***Neighborhood PC7*** | | | |
| Horvath age acceleration | **1.8 (0.4, 3.1)** | 1.3 (-1.4, 3.9) | **2.1 (0.6, 3.6)** |
| Hannaum age acceleration | **1.7 (0.4, 3.0)** | 0.9 (-1.4, 3.2) | **2.2 (0.6, 3.8)** |
| PhenoAge acceleration | **2.1 (0.4, 3.8)** | 0.8 (-2.2, 3.9) | **2.3 (0.3, 4.7)** |

[A]Models are adjusted for race/ethnicity, education level, employment, smoking status, alcohol intake, and years residing in current neighborhood
